# Leaf rolling in maize crops: from leaf scoring to canopy level measurements for phenotyping

**DOI:** 10.1101/201665

**Authors:** F. Baret, S. Madec, K. Irfan, J. Lopez, A. Comar, M. Hemmerlé, D. Dutartre, S. Praud, M. H. Tixier

## Abstract

Leaf rolling in maize crops is one of the main plant reactions to water stress that may be visually scored in the field. However, the leaf scoring did not reach the high-throughput desired by breeders for efficient phenotyping. This study investigates the relationship between leaf rolling score and the induced canopy structure changes that may be accessed by high-throughput remote sensing techniques.

Results gathered over a field phenotyping platform run in 2015 and 2016 show that leaf starts to roll for the water stressed conditions around 9:00 and reaches its maximum around 15:00. Conversely, genotypes conducted under well watered conditions do not show any significant rolling during the same day. Leaf level rolling was very strongly correlated to canopy structure changes as described by the fraction of intercepted radiation *fIPAR*_*WS*_ derived from digital hemispherical photography. The changes in *fIPAR*_*WS*_ were stronly correlated (*R*^2^=0.86, *n*=50) to the leaf level rolling visual score. Further, a very good consistency of the genotype ranking of the *fIPAR*_*WS*_ changes during the day was found (ρ=0.62). This study demonstrating the strong coordination between leaf level rolling and its impact on canopy structure changes poses the basis for new high-throughput remote sensing methods to quantify this water stress trait.

**Highligh:** The diurnal dynamics of leaf rolling scored visually is strongly related to canopy structure changes that can be documented using Digital hemispherical photography. Consequences for high-throughput field phenotyping are discussed

## 1 Introduction

Drought is recognized as one of the main factors limiting the production of maize crops (Farhangfar et al., 2015). Plants have developed several mechanisms to mitigate the impact of environmental stresses including “leaf rolling”. Under severe stress conditions, leaf lamina rolls transversally to the mid rib. This mechanism results from a differential top-bottom elastic shrinkage in the leaf cross section (Moulia, 2000). Leaf rolling has thus been related to the water potential in the leaf (Kadioglu *et al.*, 2012) and was called for this reason hydronastic (Moulia, 2000). For maize, leaf rolling is observed from leaf water potentials of -1MPa and reaches its maximum around -2MPa (Moulia, 1994). Leaf rolling reaches its maximum close to solar noon during bright sunny days (Kadioglu and Terzi, 2007) and the top leaves are generally more affected (Tatar *et al.*, 2010). This occurs when the evaporative demand is no more balanced by soil water extraction by the root system.

The leaf water potential is mostly controlled by the osmotic component through a range of biochemical pathways. Leaf rolling has been related to the accumulation of phyto-hormones (Krishna, 2003; Takahashi and Kakehi, 2010; Talaat and Shawky, 2012). Some of these hormones control stress responsive gene expression (Divi *et al.*, 2010). Some prominent changes in concentration of organic acids or ions such as K^+^ and Cl^−^ may also induce leaf rolling as demonstrated by (Saglam *et al.*, 2010). In addition to the biotic factors described earlier, (Kadioglu *et al.*, 2012) demonstrated that herbivores, viruses, bacteria and fungi may also induce leaf rolling through other biochemical pathways.

When the leaf is considered as a thin shell that verifies the law of mechanics, the transversal leaf rolling is coupled with longitudinal changes in the leaf curvature. (Hay et al., 2000; Moulia, 2000). This makes the leaf stiffer and more erect because the leaf insertion angle is generally closer to the vertical than the average leaf angle inclination. As a consequence, leaf rolling reduces the leaf surface exposed to sun light. This potentially decreases both transpiration and photosynthesis at the canopy level (Abd Allah, 2009). However stomata are generally closed under such stress conditions prevailing during leaf rolling, limiting the exchanges of CO2 and water between the leaf and the atmosphere. Nevertheless, the boundary layer resistance of the rolled leaves is increased, limiting the leaf transpiration rate (O'Toole et al., 1979). Another consequence of leaf rolling is to re-orient the normal of leaf surfaces generally away from the sun direction (Smith, 1997). This reduces the density of photon flux per unit leaf area (Duncan, 1971), limiting leaf overheating and the associated damages of the photosynthetic apparatus (Nar et al., 2009; Sarieva et al., 2010). Leaf re-orientation affects also the fraction of adaxial or abaxial faces exposed to the incoming light that have distinctive behaviors (Driscoll et al., 2006; Soares et al., 2008) with consequences on photosynthetic capacity and possible damages on the photosynthetic machinery. Leaf rolling contributes thus to maintain the internal plant water status (Subashri, 2009). On a longer time scale, leaf rolling may be also associated to a decrease in chlorophyll content due to the reduction of leaf area exposed to the sun as proposed by (Subashri, 2009) although this could also mainly result from a direct effect of drought on chlorophyll content as reported by (Bolanos and Edmeades, 1996).

Leaf rolling as a consequence of water stress results from a combination of factors including the root development, the root water extraction efficiency, the adjustment of leaf area index, the canopy structure differences under non-stressed conditions, the leaf transpiration rate and the sensitivity of the roll up mechanism to leaf water potential. Although leaf rolling observed at the canopy scale appears to be a complex trait, it bears key information on the strategy followed by the plants in case of stress conditions and should be of high value for plant breeders to evaluate genotypes. The genetic diversity in maize shows a large range of drought tolerance that is exploited by plant breeders (Adebayo and Menkir, 2014). Among several traits, leaf rolling is thus a potential trait that may be used by breeders to evaluate drought resistance. It was already associated to QTLs in rice (Price *et al.*, 2002) and durum wheat (Peleg *et al.*, 2009).

Several methods have been proposed to quantify leaf rolling. (Sirault *et al.*, 2015) evaluated the capacity of leaves to roll up under controlled conditions: leaf strips were immersed in a polyethylene glycol solution at a range of concentration. After equilibrium was reached, the leaf cross-section was imaged using micro-photographs. The convex hull of the cross section was finally exploited to quantify the leaf rolling. Several methods have been also developed to evaluate the actual level of leaf rolling under natural conditions. (O'Toole *et al.*, 1979) proposed to use a template of schematic transversal leaf sections (from flat to completely rolled position). This method was applied by (Clarke, 1986) to relate leaf rolling to leaf water concentration. A similar scoring method based on the ratio of rolled leaf width to unrolled leaf width (Premachandra *et al.*, 1993) was used to relate leaf rolling to drought tolerance (Saruhan *et al.*, 2012; Saglam *et al.*, 2014).

All the methods of leaf rolling evaluation formerly listed are relatively low-throughput. They are difficult to be applied over large phenotyping experiments because of the highly dynamic nature of leaf rolling. High-throughput leaf rolling methods are thus highly desired for field experiments. This may be completed by imaging the diurnal changes in canopy structure related to leaf rolling using high-throughput techniques based on UAV observations (Sankaran *et al.*, 2015). The objective of this study is to quantify the effect of leaf rolling on canopy structure using Digital Hemispherical Photograph (DHP).

DHP provides a very efficient way to describe several canopy structure variables from the directional gap fraction Po(θ) (Jonckheere *et al.*, 2004; Weiss *et al.*, 2004b; Lopez-Lozano *et al.*, 2007). A total of 50 maize genotypes with contrasting leaf rolling behavior was studied during two consecutive years. The diurnal variation of canopy structure of each genotype was documented using DHP measurements completed several times during the day under severe water stress conditions. The consistency of DHP measurements with visual scoring of leaf rolling was investigated. A method is then proposed to quantify the leaf rolling from the DHP. Results are finally discussed on the possible use of UAV observations for high-throughput leaf rolling characterization.

## 2 Materials and methods

### 2.1 The experiment

Two experiments were conducted near Nérac, France (44.17° N, 0.30° E) in 2015 and 2016. The rows were oriented NW-SW. A total of 800 genotypes of maize were grown in plots made of 2 adjacent rows with 0.8m spacing by 6 m long (Figure 2). A subsample of 38 genotypes were selected for their large differences in canopy structure and susceptibility to leaf rolling. In 2015, 30 genotypes were maintained under severe water stress conditions (WS modality). In 2016 16 genotypes were maintained similarly under severe water stress conditions (WS modality) while 4 of them were also conducted under well irrigated conditions (WW modality). Height genotypes were present on both years in the WS modality while only 2 of them were also sampled in WD modality in 2016 (Table 1). The soil moisture at the field capacity is 200 mm, with hardly available water (HAW) below 60 mm. In 2015, the soil moisture was below HAW since the 5^th^ of July for the WS modality, with 25 mm remaining water the 5^th^ of August at the date of the leaf rolling measurements. In 2016, the soil moisture was below HAW since the 8^th^ of July, with 35 mm remaining water the 3^rd^ of August at the date of the leaf rolling measurements. Regarding the soil conditions, the water stress was therefore slightly stronger in 2015 as compared to 2016. The leaf rolling measurements were performed roughly at the female flowering stage known to be very sensitive to water stress. For both years, measurements were completed under very hot and sunny days with almost the same illumination conditions (Figure 1). However, the 5^th^ of August 2015 was more stressful as compared to the 3^rd^ of August 2016, with higher temperatures and much larger vapor pressure deficit (VPD) as calculated from air temperature and humidity after (Monteith and Unsworth, 2007).

**Figure 1.**
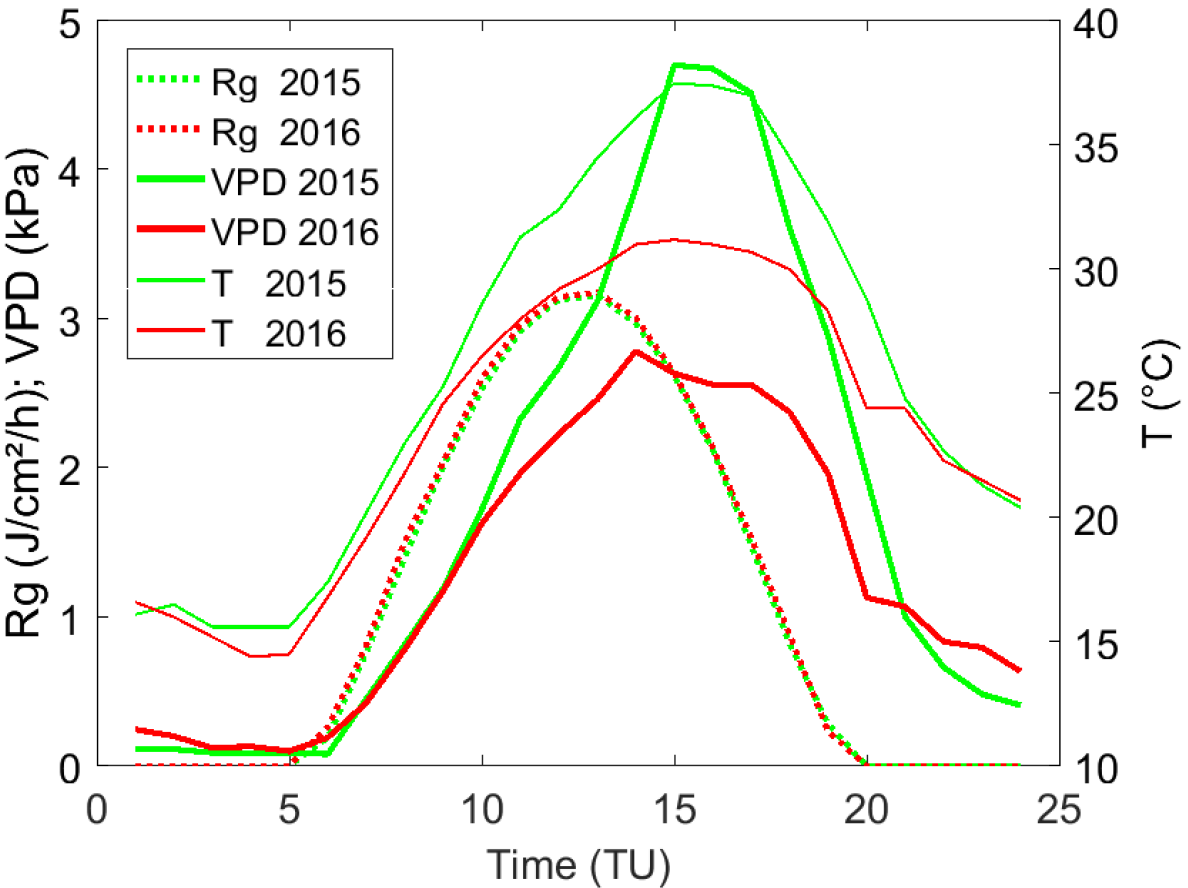
diurnal variation of global radiation (Rg), temperature (T) and VPD observed the 5^th^ of August 2015 and the 3^rd^ of August 2016 in Nérac experiment.

**Figure 2:**
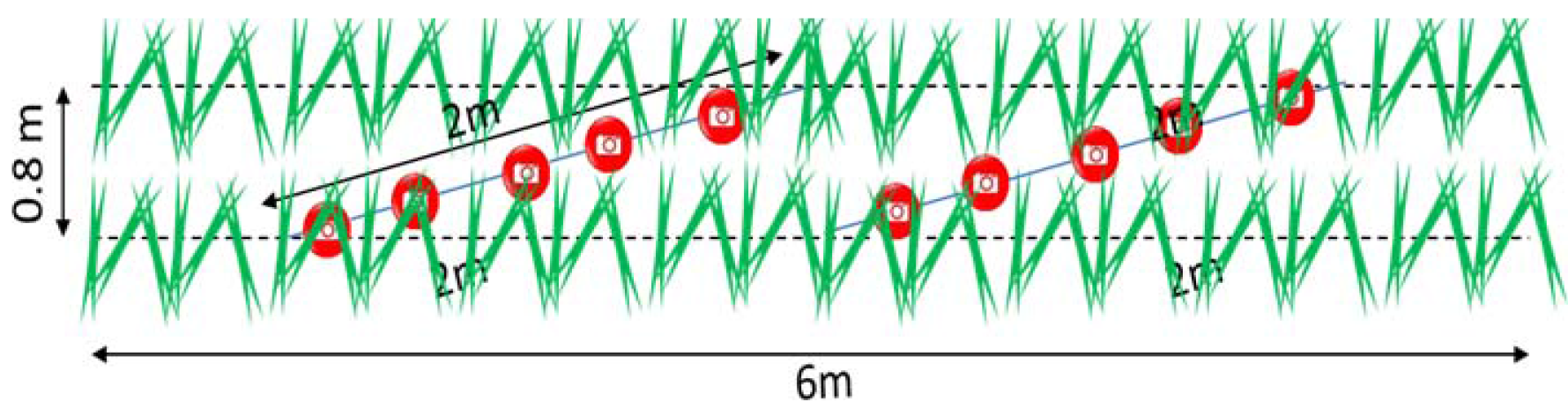
Location of DHP measurements over the 2 rows of a microplot.

**Table 1.** Distribution of the 38 genotypes used in 2015 and 2016 for the WS and WD modalities.

| **Genotypes #** | **WS 2015** | **WS 2016** | **WW 2016** | **Number** |
| --- | --- | --- | --- | --- |
| 16-37 | X |  |  | 22 |
| 10-15 |  | X |  | 6 |
| 2, 4, 5, 6, 7, 8 | X | X |  | 6 |
| 0, 1 |  | X | X | 2 |
| 3, 9 | X | X | X | 2 |
| **Total** | **30** | **16** | **4** | **50** |

### 2.2 Visual scoring of leaf rolling

The leaf rolling was scored by some visual notations from 1 to 9: score 1 corresponds to no leaf rolling (the cross section of the leaf is almost flat) which observed during early morning or under no water stress conditions; score of 9 corresponds to the maximum leaf rolling, i.e. when the leaf cross section is fully rolled. The same operator was scoring the microplots for the two experiments, to limit possible biases during the day and years. The scoring was repeated approximately every hour from 9:00 to 17:00. It results in a total of 430 notations. About 15 minutes were necessary to score 30 plots.

### 2.3 DHP measurements

Upward looking digital hemispherical photographs were taken with a sigma SD-14 equipped with fisheye lens of 8 mm focal length. The camera was set on automatic exposure. The images are recorded in jpg with 2640 × 1760 pixels. The optical center and projection function of the camera were calibrated using the method described in (http://www.avignon.inra.fr/can_eye). Images were repeated approximately every 1.5 hour during the day from 9:00 to 17:00 resulting in 7 series of 30 (in 2015) or 20 (in 2016) images. About 30 minutes were necessary to sample 30 plots.

A total of 10 photos were taken on each microplot at each time step in the day to capture the spatial variability (Weiss *et al.*, 2004a). Photos were distributed over two diagonal segments placed between the two center rows (Figure 2). A 2m long stick with marked positions was used to indicate the precise location of the camera for taking each image over a segment. The position of the stick over each microplot was kept the same across the repeated measurements during the day to provide a high degree of temporal consistency. The camera was looking upward and always oriented the same with regards to the row direction. Sampling a microplot with 10 images takes about 1 minute. Note that the hemispherical images account partly for the neighboring microplots. However, this influence is limited since images are exploited for zenith angles smaller than 60°.

### 2.4 Processing the DHP

The 3500 images (50 microplots with 10 images per microplot sampled 7 times during the day) were processed using the CAN-EYE freeware (www.avignon.inra.fr/can_eye). CAN-EYE is a package of Matlab functions developed to estimate canopy structure characteristics from RGB images (Demarez *et al.*, 2008). Direct exposure of sun over the camera during afternoon measurement induces local artifacts (see S4 and S7 in Figure 3) that were easy to correct using the versatile color segmentation in CAN-EYE.

**Figure 3:**
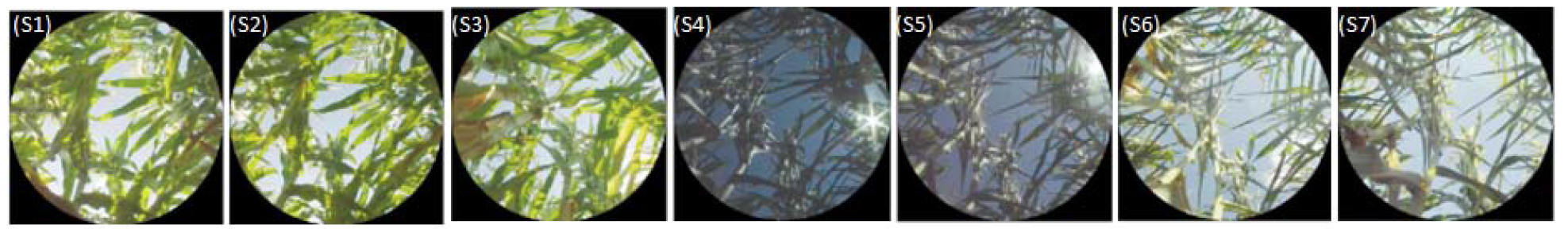
DHP images taken at seven different time (S1: morning, S7: late afternoon) at the same location of a microplot. Images show artifacts for S4 and S5 due to direct sun light. It shows clearly the changes of canopy structure from the morning to afternoon due to leaf rolling

The main output of can-eye is the bidirectional gap fraction P_*o*_(θ, φ) where 0° < θ < 62.5° and 0° < *ϕ* < 360°. θ = 0° corresponds to the nadir direction and *ϕ*=0° corresponds to the row direction. The zenith and azimuth directions are integrated into 2.5° steps. Zenith angles higher than 62.6° were discarded due to the large fraction of mixed pixel and the larger contribution of the neighboring plots. Due to the assumed symmetry along the row direction and across the row directions, the directional gap fraction values were averaged to provide a representative quadrant, 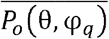 with 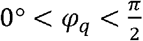:

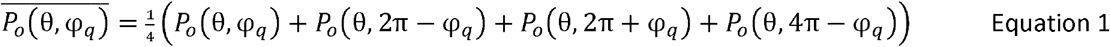

Several “segmental gap fraction” were computed to investigate the possible correlation with the leaf rolling score. They correspond to integration over larger solid angles to provide more stable directional gap fraction values: the hemisphere was divided into three different rings of 20° zenithal sectors: [0°< θ <20°], [20°< θ <40°], [40°< θ <60°]. Further, each ring was divided into three azimuthal ranges: [0°< φ_*q*_ <30°], corresponding to the row direction, [30°< φ_*q*_ <60°] corresponding to a direction diagonal to the row direction, and [60°< φ_*q*_ <90°] corresponding to the direction perpendicular to the row. Therefore, a total of 9 integrated gap fractions were computed. In addition, the fraction of diffuse radiation intercepted by the canopy, called white sky flPAR (*fIPAR*^*WS*^), was also computed:

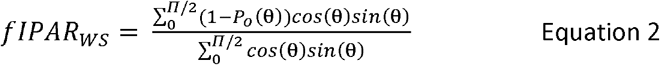

Where *P*_*o*_(θ) is the azimuthally averaged value of *P*_*o*_(θ, φ). When evaluating *flPAR*_*WS*_, values of *P*_o_(θ) for θ > 62.5° were computed assuming a linear interpolation of the term (1 − *P*_*o*_(θ))*cos*(θ)*sin*(θ) between θ = 62.5° when *P*_*o*_(62.5°) is measured, and 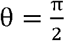 for which 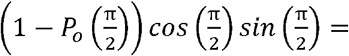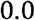.

## 3 Results and Discussion

### 3.1 Diurnal variation of the leaf rolling visual score

In absence of water stress, leaves keep unrolled as observed in Figure 1 for the 4 genotypes grown under well irrigated conditions in 2016. Conversely, leaf rolling was observed for all the genotypes subjected to water stress both in 2015 and 2016. All the genotypes show very similar diurnal patterns of the leaf rolling score both in 2015 and 2016. The score starts from the minimum value, Score≈1, in the early morning (7:30 UT) when no leaf rolling is observed (Figure 4). However, few cultivars show already some leaf rolling for the first scoring of the day. Then, leaves roll up progressively when the water stress experienced by the canopy increases as a function of the climatic demand controlled mainly by the incoming radiation, and the vapor pressure deficit: at 9:00 UT leaf rolling was already observed on many genotypes on both years, when VPD≈1.5 kPa. Maximum leaf rolling was reached around 15:30 UT with some significant variation of the magnitude between genotypes. This corresponds to the maximum value of the VPD during the day (Figure 1). Finally, leaves start to unroll when the climatic demand decreases significantly. Variability at a given time between genotypes under water stress is maximum when the rate of increase of leaf rolling score is maximum, around 12:30 UT (Figure 4).

**Figure 4:**
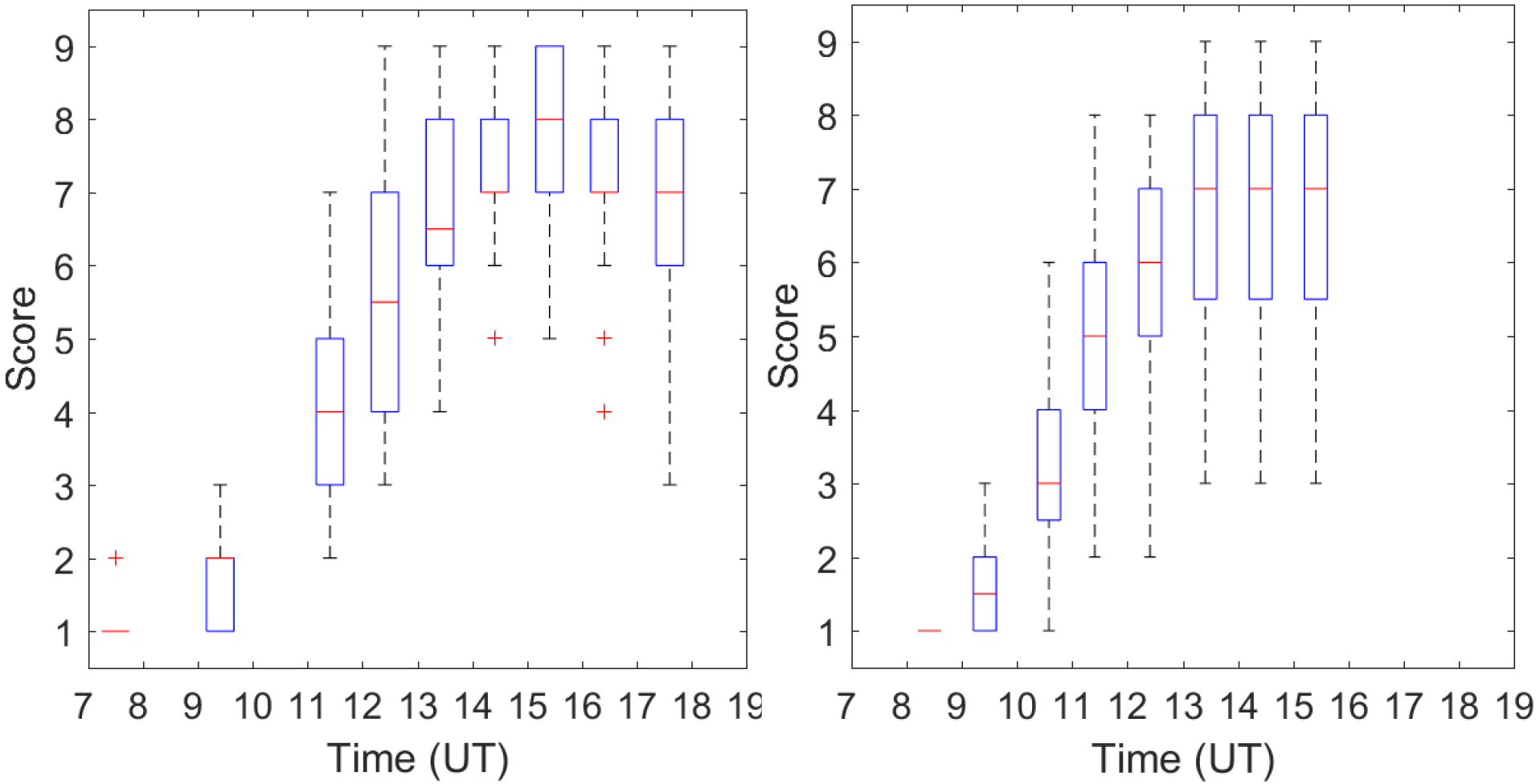
Diurnal pattern of leaf rolling scores for Water stress modality. In 2015 (left) 30 genotypes; in 2016 (right) 16 genotypes. Box plot representation where the red line is the median, the edges of the box are the 25th and 75th percentiles, the whiskers extend to the most extreme data points the algorithm considers to be not outliers, and the outliers are plotted individually as red ‘+’.

### 3.2 Diurnal variation of the directional gap fraction

Similarly to the visual score, no clear diurnal variation of the gap fraction is observed over the irrigated genotypes (Figure 5, red curves on the right). Conversely, the directional gap fraction increases during the day over the water stressed genotypes both in 2015 and 2016 (Figure 5, black curves). Very similar patterns to those of leaf rolling scores are observed, with a minimum value in the early morning, and a maximum value reached around 15:30 UT. This corresponds to the maximum daily temperature and VPD values (Figure 1). After this maximum value, leaves start to unroll at the end of the afternoon.

**Figure 5:**
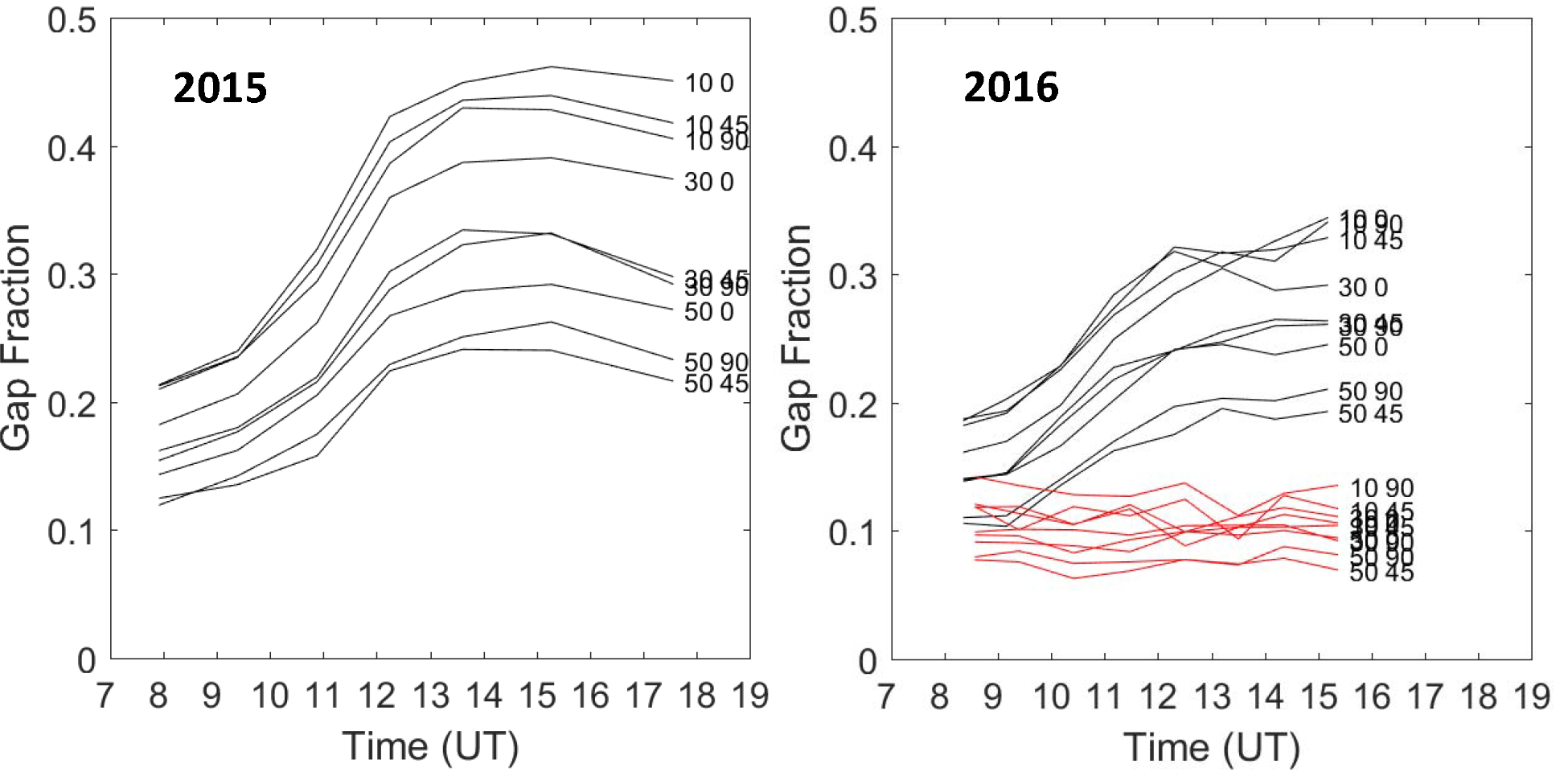
Diurnal evolution of the gap fraction values measured for 9 solid angles indicated by the numbers on the left: the first number if the zenith, the second one is the azimuth. Each curve is the average of the 30 genotypes for 2015, 16 water-stressed genotypes for 2016, and 4 irrigated genotypes for 2016.

The same diurnal pattern is observed for all the directions considered, with mainly changes in minimum and maximum values (Figure 5). As expected, higher gap fraction values are observed close to nadir and in the directions parallel to the row. Conversely, the lowest gap fraction values are observed for directions perpendicular to the row and for the larger zenith angles. If the largest diurnal variation is observed for the near nadir directions, a very strong consistency is observed between the diurnal patterns of all the directions considered (Figure 5).

### 3.3 Impact between leaf level rolling and canopy architecture changes

The leaf level rolling as scored visually was tentatively related to the directional gap fractions measured with the DHPs that document the corresponding changes in canopy architecture. The fraction of intercepted radiation under diffuse illumination conditions (*fIPAR*_*WS*_) was proposed to be used as a proxy of the canopy structure. This is supported by the very strong consistency between the gap fractions observed in the different directions (Figure 5). The *flPAR*_*WS*_ is computed from the directional integration of the gap fractions (Equation 2). The integration offers the advantage to smooth out uncertainties associated to each directional gap fraction measurements, providing therefore more robust results.

The visual score corresponding to the unrolled state of the leaf observed in the early morning (before 9:00 UT) were always set to Score≈1 for each genotype (Figure 4). However, at the canopy level, *fIPAR*_*WS*_ values show significant variability between genotypes when leaf rolling has not yet started (Figure 6). These differences are explained by genotypic specificities in the canopy architecture due to differences in the leaf area index and/or in plant morphology. Note that the irrigated modality (WW) in 2016 shows the higher *fIPAR*_*WS*_ values as compared to the water stress (WS) modalities since drought already impacted leaf expansion during the weeks preceding the measurements. Similarly, the 2016 water stress was less severe as in 2015, with generally larger *fIPAR*_*WS*_ values in 2016, in agreement with the water balance presented previously. Closer inspection of the distribution of the 8 genotypes that are common between 2015 and 2016 experiments show that the values of the unrolled *flPAR*_*WS*_ observed in 2015 are not correlated with those observed in 2016 (R^2^=0.07) with a very small spearman correlation coefficient (ρ^2^=0.06).

**Figure 6.**
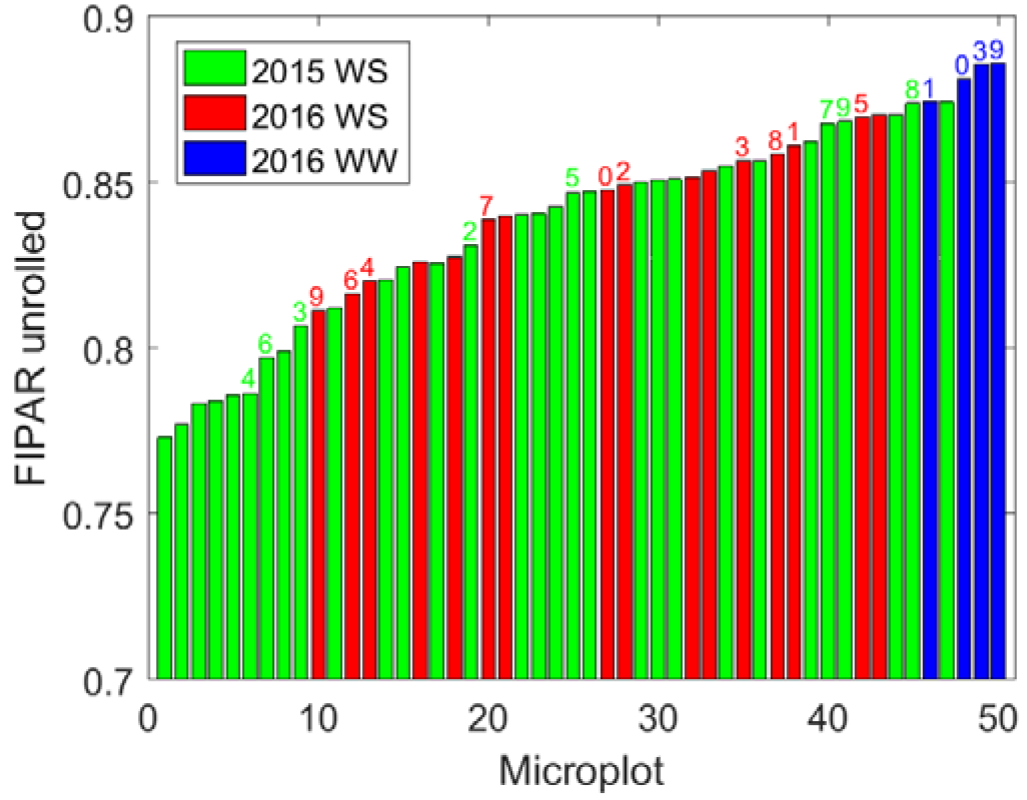
Distribution of the *fIPAR*^*WS*^ values observed in the early morning (unrolled state) for all the 50 genotypes investigated. The values are sorted in ascending order. The colors correspond to the years and modalities. The genotypes common between years and modalities are indicated above each bar.

The differences in *fIPAR*_*WS*_ values in the early morning between genotypes and environmental conditions induced differences in the *flPAR*_*WS*_ values observed during maximum leaf rolling as demonstrated by Figure 7, left panel (R^2^=0.50, n=46, correlation significant at α=5%). However, no significant (α=5%) correlation is observed between the difference 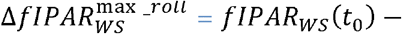 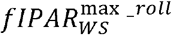 between the unrolled state in the early morning, *fIPAR*_*WS*_(*t*_0_), and the minimum *fIPAR*_*WS*_ value corresponding to maximum leaf rolling in the late afternoon, 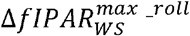 (Figure 7, right panel). A simple base line normalization is therefore proposed to limit the impact of the genotypic and environmental differences in the early morning: the *fIPAR*_*WS*_(*t*) values observed during the day at time *t* are subtracted from the unrolled *fIPAR*_*WS*_(*t*_0_) values observed in the early morning, *t*_0_.: Δ*fIPAR*_*WS*_ = *fIPAR*_*WS*_(*t*_0_) − (*t*)

**Figure 7.**
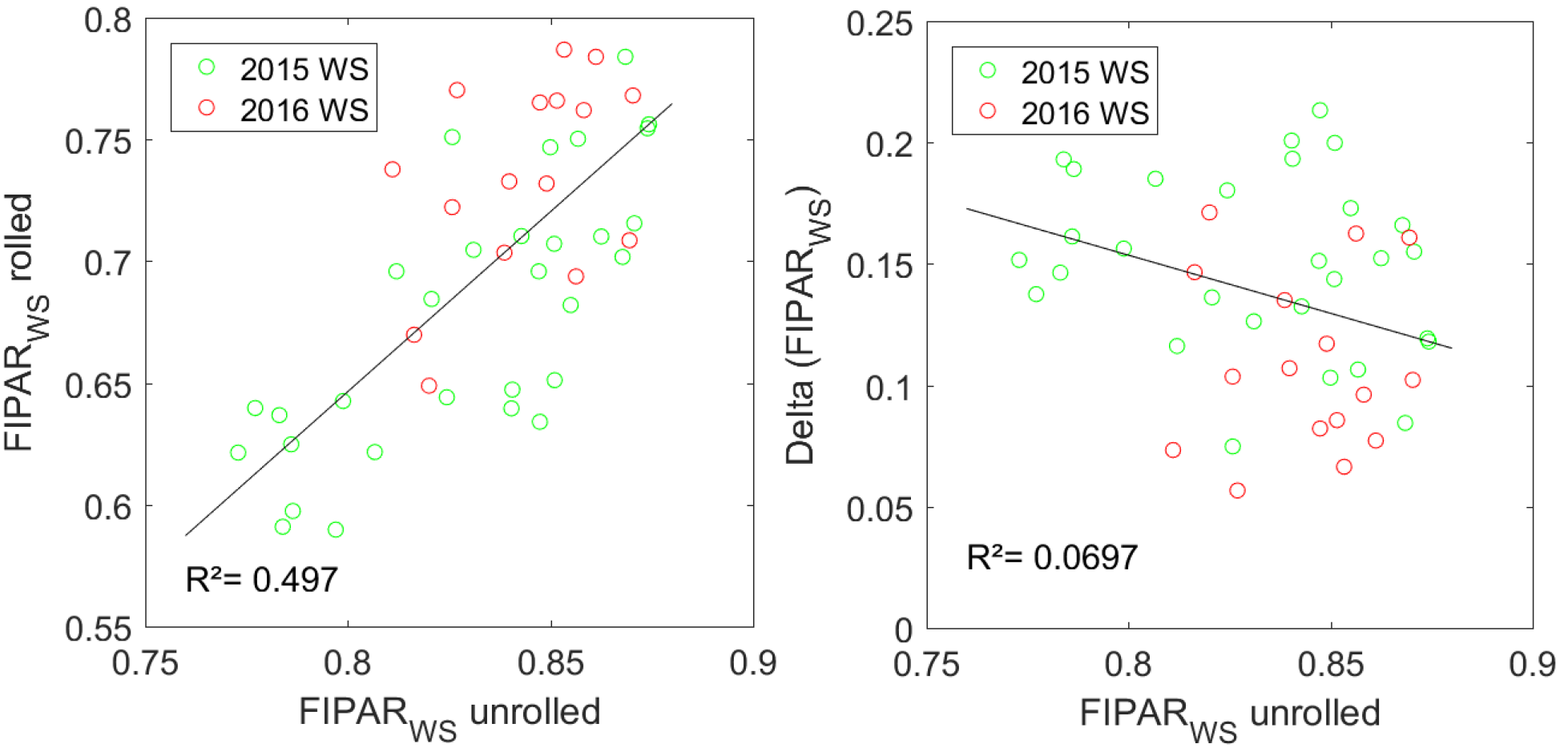
On the left panel, relationship between the unrolled state *fIPAR*_*WS*_ observed in the early morning (FIPARWS unrolled) and the *fIPAR*_*WS*_ values corresponding to the state with maximum leaf rolling observed in the late afternoon (FIPAR_WS_ rolled). On the right panel, relationship between the unrolled state *fIPAR*_*WS*_ (FIPAR_WS_ unrolled) and the difference between *fIPAR*^*ws*^ values observed between early morning and late afternoon (Delta (FIPAR_WS_)). Data corresponding to the water stress modality (WS, 46 points) in 2015 (green) and 2016 (red). The solid line represents the linear best fit.

The visual scores and the corresponding *fIPAR*_*WS*_(*t*) values need to be assigned to the same time during the day to establish a relationship between them. The visual scores that were more frequently sampled were thus linearly interpolated at the time of the DHP measurements from which *fIPAR*_*WS*_(*t*) were computed.

Results (Figure 8) show that the constraint in the early morning between the leaf rolling score (*Score*(*t*_0_) = 1.0) and the Δ*fIPAR*_*WS*_ values is well verified later in the day when no leaf rolling is experienced such as for the irrigated modality in 2016. The Δ*fIPAR*_*WS*_ values are strongly and linearly related to the *Score* (Figure 8). A simple linear model verifying the early morning constraint (*Score*(*t*_0_) = 1.0; Δ*fIPAR*_*WS*_(*t*_0_) = 0.0) was fitted to the available data:

**Figure 8.**
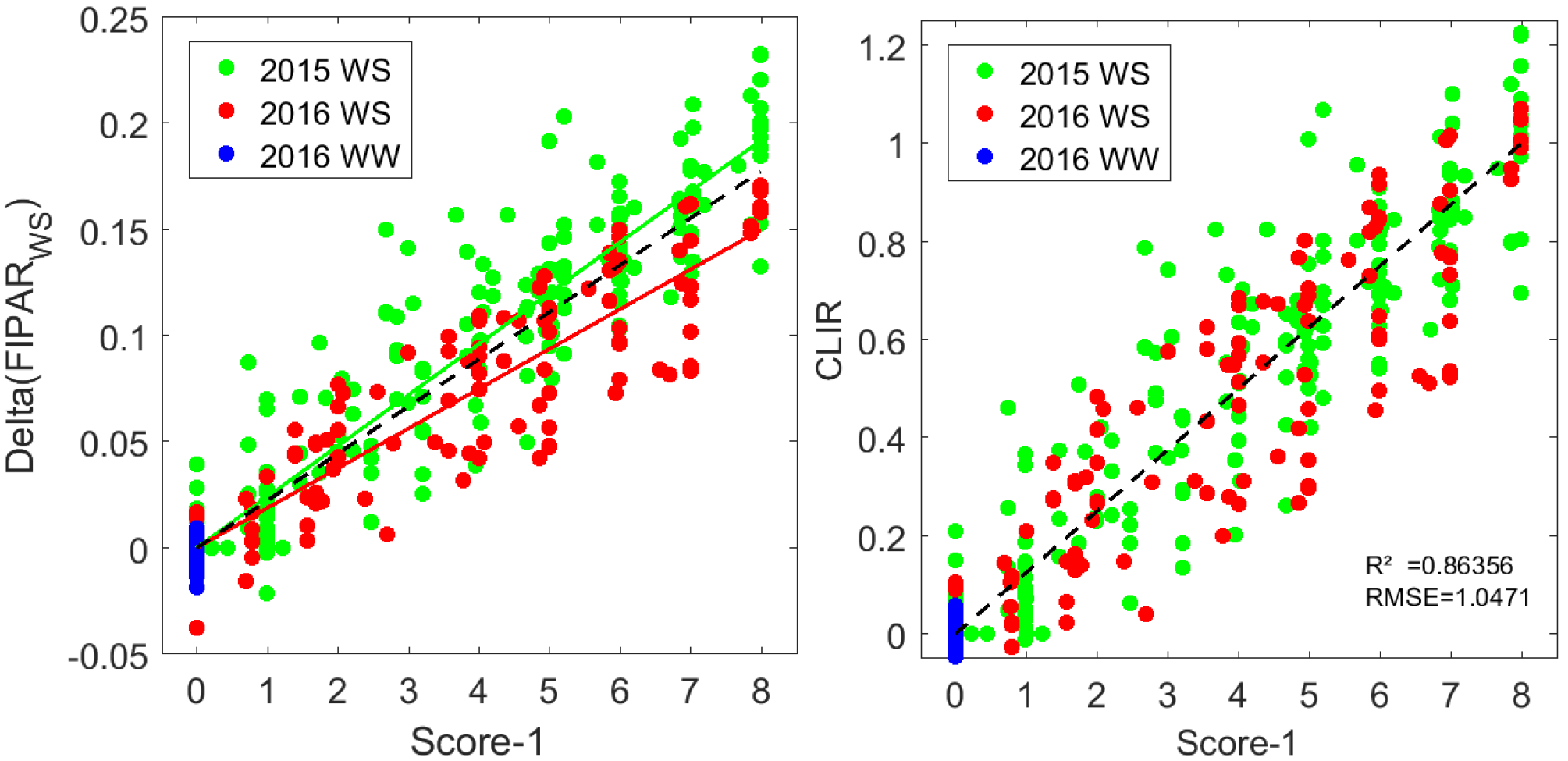
On the left, relationship between Δ*fIPAR*_*WS*_ and the leaf rolling visual score (Score-1). The solid lines correspond to the best fit line verifying the constraint Δ*fIPAR*_*WS*_ = 0 when *Score* = 1 (Equation 3) for the 2015 (green) and 2016 (red) WS modalities. The black dash line corresponds to the best fit over the 370 available points (including the WW modality in 2016). On thr right, relationship between CLIR and the leaf rolling visual score (Score-1). The solid black line corresponds to Equation 5.

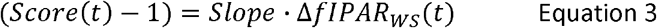

It provides very good performances (Table 2) for the 2 years under water stress conditions. For the irrigated modality in 2016, the points are concentrated close to the unrolled leaf situation with *Score*(*t*_0_) = 1.0 and Δ*fIPAR*_*WS*_(*t*_0_) = 0.0. However, a small difference is observed between the two years, 2016 showing lower sensitivity of the canopy structure (Δ*fIPAR*_*WS*_) to the leaf level rolling (*Score*). This difference may partly be attributed to the slightly smaller leaf development of the canopy in 2016 as observed in Figure 6: when the canopy is less developed, limited absolute effects on canopy structure are expected. This effect may be accounted for by normalizing the values of Δ*fIPAR*_*WS*_ by the average Δ*fIPAR*_*WS*_ value observed for the maximum leaf rolling state, 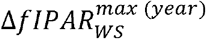, leading to propose the *CLIR* index (Canopy Level Index for Rolling):

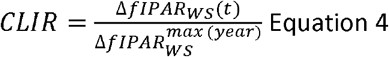

**Table 2.** Characteristics of the models (Equation 3 or Equation 4) used to relate the leaf rolling visual score to the canopy *fIPAR*_*WS*_ level values. RMSE is expressed in Score units.^(1)^ It includes also the 2016 WW plots.

| Model | Sample | Nb. points | Slope | R<sup>2</sup> | RMSE |
| --- | --- | --- | --- | --- | --- |
| $(Score(t) - 1) = Slope \cdot \Delta fIPAR_{WS}(t)$ | 2015 WS | 210 | 41.64 | 0.857 | 1.05 |
|  | 2016 WS | 128 | 53.32 | 0.823 | 1.17 |
|  | All<sup>(1)</sup> | 370 | 45.04 | 0.845 | 1.14 |
| $(Score(t) - 1) = 8 \cdot CLIR(t)$ | All<sup>(1)</sup> | 370 | 8 | 0.864 | 1.05 |
| $(\max(Score - 1)) = 8 \cdot \max(CLIR)$ | All<sup>(1)</sup> | 50 | 8 | 0.771 | 1.15 |

The canopy level leaf rolling index, *CLIR*, is theefore related to the leaf level rolling score according to:

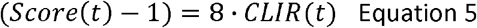

Where 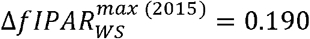 and 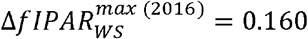. Results (Table 2) show a slight improvement in the performances of the regression due to the enhanced consistency between years.

### 3.4 Comparison between genotypes

The comparison of the reaction to water stress between genotypes was completed by considering either the magnitude of the diurnal variation of leaf rolling or the way it develops, i.e. the dynamics.

#### 3.4.1 Magnitude of leaf rolling

The magnitude of the leaf rolling score was evaluated as max (*Score* − 1) over the day for each microplot. The well-watered modality in 2016 (2016 WW) have a minimal magnitude consistently with the no leaf rolling experienced. Under water stress conditions, genotypes show important differences (Figure 9 left). About half the microplots have a maximum rolling score lower or equal to 6, with few of them showing only small leaf rolling at the end of the afternoon when it is expected to be maximum. Year 2016 appears to have generally less leaf rolling in agreement with the previous observations (Figure 4). The 8 genotypes common in 2015 and 2016 show relatively low level of consistency of the ranking between the 2 years (Figure 10 left). Conversely, a high degree of consistency is observed when using the CLIR index to quantify leaf rolling at the canopy level (Figure 10 right). The better performances for CLIR as compared to the leaf Score are mainly explained by the more progressive values of *CLIR* as observed on the distribution of values (Figure 9 right) as well as the more objective values provided by the DHP measurements as compared to the visual scoring. Nevertheless the two quantities are highly correlated (Table 3) with R^2^=0.771 and RMSE=1.15.

**Figure 9.**
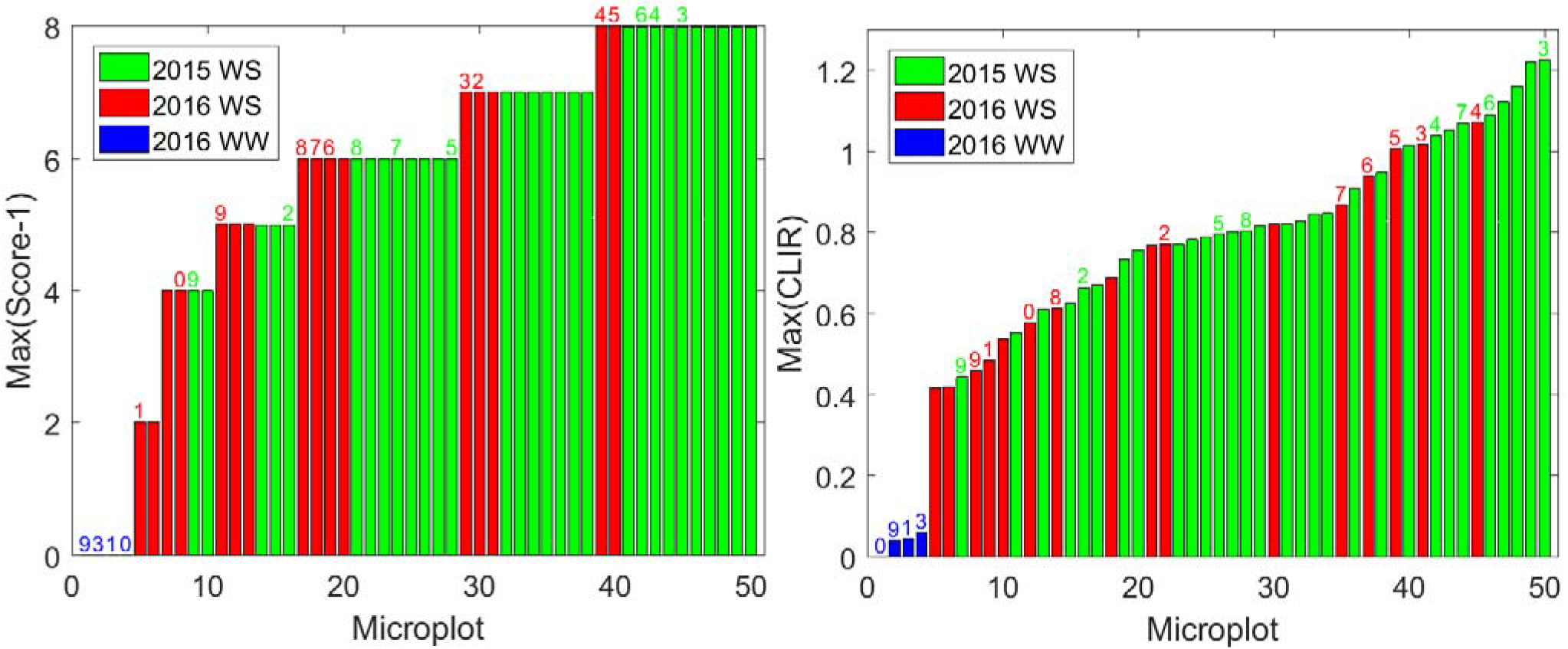
On the left, distribution of the maximum value of the leaf rolling visual score(Max(Score-l)) observed over each microplot during the day. On the right, distribution of the maximum value of *CLIR* observed over each microplot during the day. The values are sorted in ascending order. The colors correspond to the years and modalities for the 50 microplots. The genotypes common between years and modalities are indicated above each bar.

**Figure 10.**
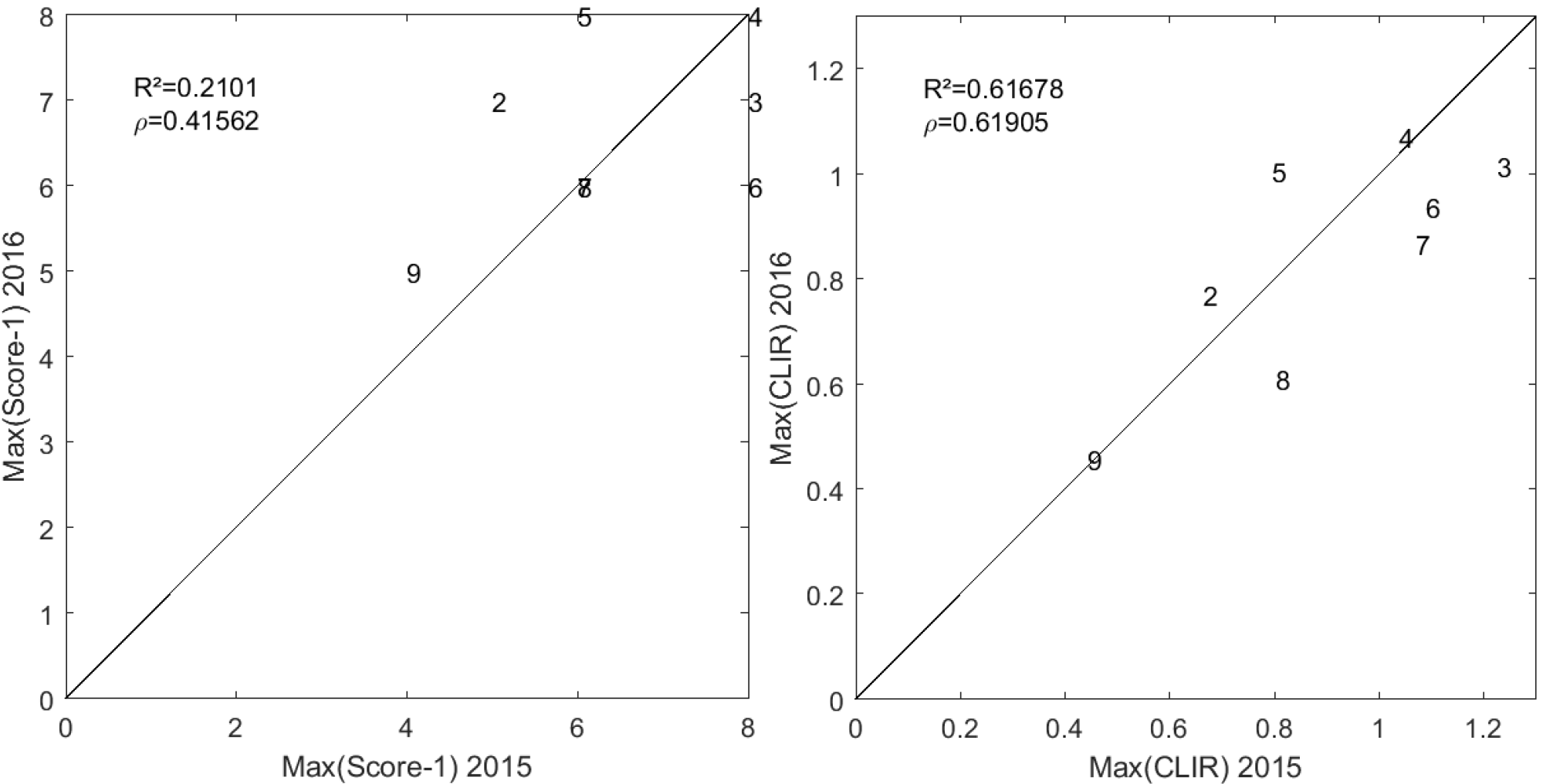
On the left, comparison between the maximum value of the normalized score (Score-1) observed in 2015 (x axis) and 2016 (y axis). On the right, comparison between the maximum value of *CLIR* observed in 2015 (x axis) and 2016 (y axis). The numbers correspond to the genotype identifiant. The solid line is the 1:1 line. The Pearson (R^2^) and Spearman (ρ^2^) coefficients are provided.

#### 3.4.2 Diurnal dynamics of the leaf rolling

The effect of leaf rolling at the canopy level was investigated here by transforming the *fIPAR*_*WS*_ values into leaf rolling score according to equation 4. The diurnal dynamics of the leaf rolling visual score and the estimates from the DHP measurements are generally very consistent (Figure 11), confirming the previous results (Table 1 and Figure 8). The well-watered modality shows no leaf rolling at all along with no significant changes in the canopy structure during the day (Figure 11).

**Figure 11.**
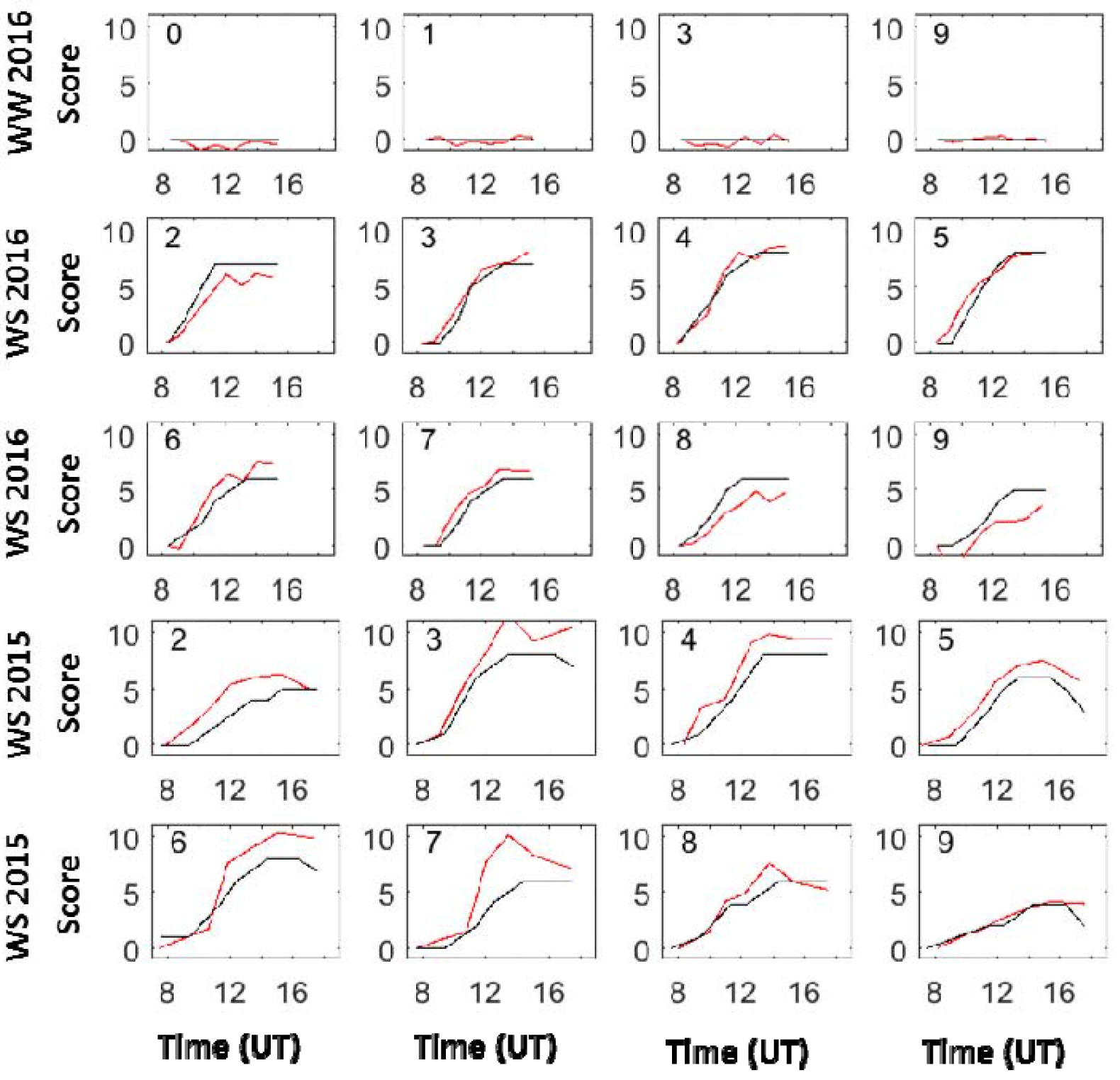
Diurnal dynamics of the leaf rolling visual scores evaluated at the leaf level (black line), and the score estimated from equation 5 (red line) from canopy level DHP measurements. Top, the 2 center and bottom plots correspond respectively to 2016 WW (4 genotypes), 2016 WS (8 genotypes) and 2015 WS (the same 8 genotypes). Genotype identifiant is given in each subplot.

Although the genotypic variability of the magnitude of rolling at the leaf and canopy levels appears as one of the main features observed, other traits of the dynamics were further investigated. For this purpose, both the visual scores at the leaf level and the canopy level *CLIR* values derived from the DHP measurements were normalized by their minimum and maximum values observed in the 8:00 to 17:00 time interval over each plot:

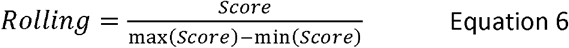

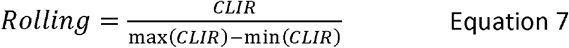

Emphasis was then put on the period of rapid rolling variation corresponding to the 9:00 to 13:00 time interval (Figure 11). The very good consistency between both types of measurements is further demonstrated by Figure 12. The development of leaf and canopy level rolling appears very linear with time. Two traits characterizing the dynamics were therefore computed for each plot using a robust linear fit to the available data: the slope (*α*) and the time at half maximum (*t*_*max*/2_).

**Figure 12.**
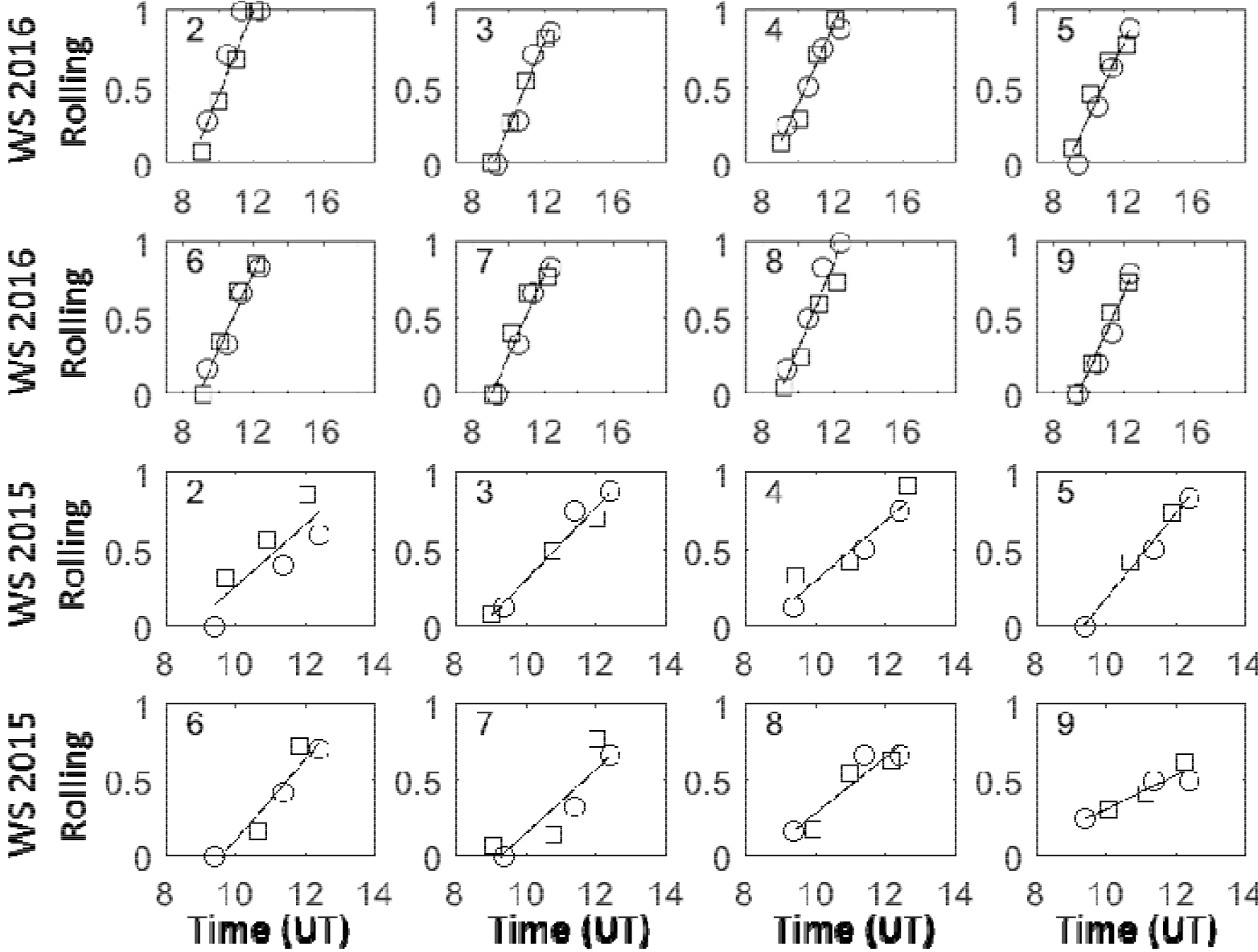
dynamics of the rolling evaluated at the leaf level (O), and the canopy level (⸀) using Equation in the 9:00 to 13:00 time interval. The solid line corresponds to the best linear robust fit. The 2 top and bottom plots correspond respectively to 2016 WS (8 genotypes) and 2015 WS (the same 8 genotypes). Genotype identifiant is given in each subplot.

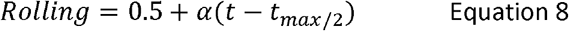

Results show that *t*_*max*/2_ varies strongly between genotypes and years: 11:15 < *t*_*max*/2_ < 12:45 (Figure 13 left). Earlier *t*_*max*/2_ is observed in 2016 as compared to 2015. Consistency between the 8 common genotypes across the 2 years is relatively poor (ρ=0.33, R^2^=0.49). The slope (α) shows also a large variability between genotypes and years: 0.1 < ∝ < 0.3 (Figure 13 right and Figure 14 right). The consistency between the 8 common genotypes among the 2 years for α values is even poorer (ρ=0.29, R^2^=0.21) than that of *t*_*max*/2_. Conversely to *t*_*max*/2_, larger values of ∝ are generally observed in 2016 (Figure 13 right and Figure 14 left). A negative correlation links *t*_*max*/2_ and α (R^2^=0.42): since rolling starts to develop approximately at the same time (between 8:15 to 9:30, Figure 11 and Figure 12), the value of the half magnitude will be reached for earlier *t*_*max*/2_ values. However, the values are much similar for the 2 years when computing for each plot the absolute value of the rolling development rate: while the leaf score (∝ (max(*Score*) — min(*Score*)) shows still some year effect (Figure 14 center), the canopy level (*α*(max(*CLIR*) — mim(*CLIR*)) shows marginal effects, particularly for the 8 common genotypes (Figure 14 right).

**Figure 13.**
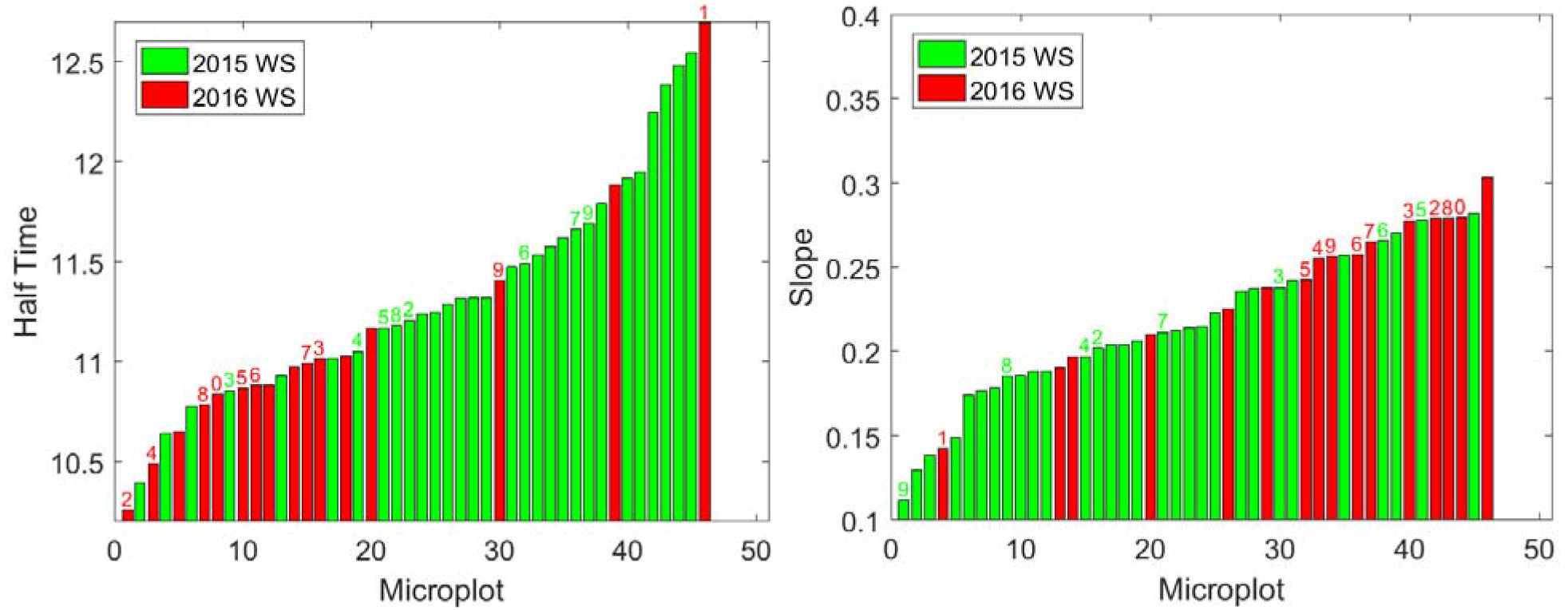
Distribution of the time (*t*_*max*/2_ in hour) when half the maximum rolling is observed (left) and the slope corresponding to the rate of change of the rolling from minimum to maximum (α in hour^−1^). The colors correspond to the years and modalities for the 46 microplots (the WW 2016 plots are obviously not represented here). The genotypes common between years and modalities are indicated above each bar.

**Figure 14.**
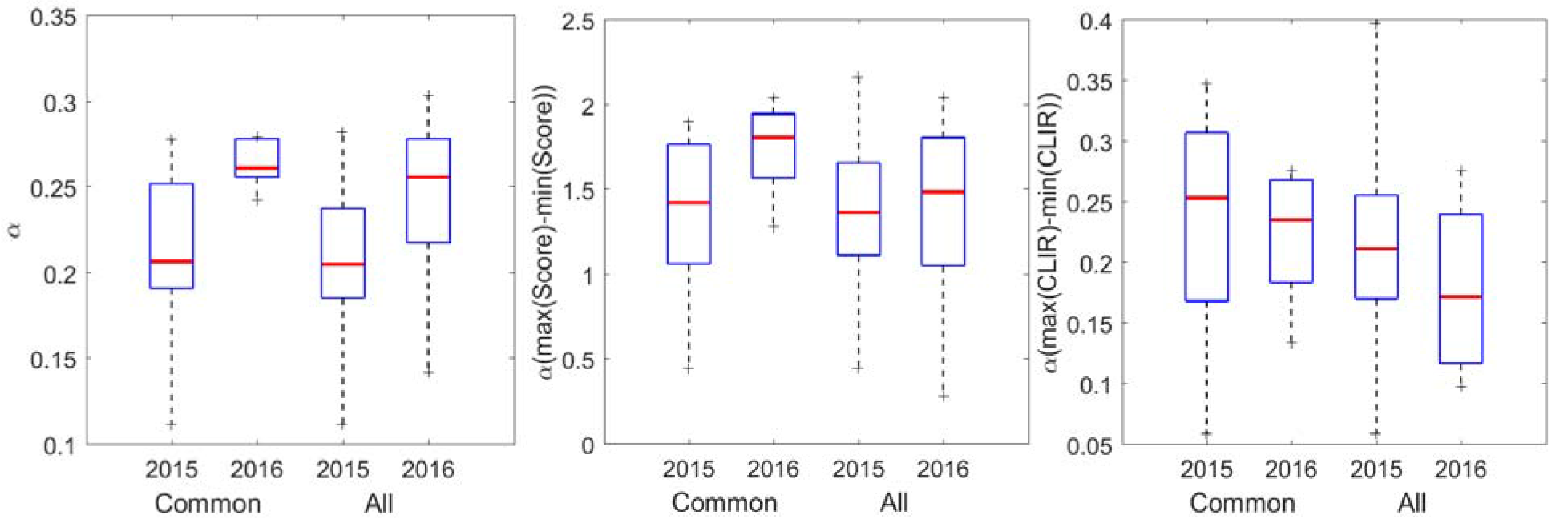
Distribution of the slope values for the 8 common genotypes in 2015 or 2016 and for all the genotypes considered in 2015 (30) and 2016 (16) for the WS modality. Three slopes are displayed: on the left the slope α computed on the normalized Scores and CLIR data; in the center, the slopes in absolute values of Scores (∝ (max(*Score*) - min(*Score*))); on the right, the slopes in absolute values of CLIR (α(max(CLIR) - min(CLIR))).

## 4 Conclusion

This study focuses on leaf rolling in maize crops subjected to water stress. During the 2 year experiments conducted around the female flowering stage, the soil moisture was below the hardly extractible water threshold, and high VPD values were observed during the day. As a consequence, leaf rolling was starting to be significant after 9:00 UT when VPD≈1.5 kPa, reaching its maximum value around 15:30 UT close to the maximum VPD value of the day. Differences between genotypes and years were mostly related to the maximum of leaf rolling score observed. Year 2016 was characterized by lower score values in relation with the smaller water stress level experienced (more water in the soil, lower VPD values) as compared to year 2015. However, the 8 genotypes sampled on both years show a relatively good consistency between both years in the ranking of the maximum score values observed (ρ=0.41) which would indicate some useful degree of heritability. Further works conducted over the ensemble of microplots of the experiments should be undertaken to confirm the level of heritability of the leaf rolling. However, scoring the leaf rolling from visual inspection of hundreds of microplots within a limited time period is not easily feasible because of the highly dynamic character of leaf rolling. Alternative high-throughput phenotyping methods are thus highly desired. This is the reason why this study investigated concurrently the impact of leaf rolling on canopy structure that may be accessed with high-throughput using remote sensing techniques. However, before developing an operational system, we concentrated on the comparison between leaf level and canopy level rolling features.

Canopy structure changes due to leaf rolling were documented using DHP measurements. More detailed inspection of the relationship between the changes in the gap fractions derived from DHPs and the leaf level visual scoring of rolling shows strong correlations for all the directions considered. For this reason, the white sky FIPAR, *FIPAR*_*WS*_, was proposed as a good proxy of the canopy structure changes induced by the leaf level rolling: it retains the main changes while smoothing out uncertainties associated with limited directional sampling of the gap fraction. To compensate for the possible differences of *FIPAR*_*WS*_ between experimental conditions and genotypes in the early morning when no leaf rolling is expected, the early morning *FIPAR*_*WS*_ values were subtracted from the *FIPAR*_*WS*_ values measured during the day to get Δ*FIPAR*_*WS*_. However, the values of Δ*FIPAR*_*WS*_ observed around 15:00 UTC when the maximum leaf rolling is expected may differ from year to year depending on the experimental conditions. This effect was further accounted for by normalizing Δ*FIPAR*_*WS*_ by the mean of maximum values of Δ*FIPAR*_*WS*_ observed during the day across all the micro-plots available. This resulted in the Canopy Level Index for Rolling (*CLIR*). The coordination between the rolling of the leaf section and the opening of the canopy as quantified by *CLIR* appears very strong and relatively stable across genotypes and the 2 years investigated. Further, a higher degree of consistency (ρ=0.62) was observed between the ranking of the maximum CLIR values of 2015 and 2016 years among the 8 genotypes common for the 2 years.

Apart from the magnitude, other features associated to the dynamics were investigated. The time when half magnitude is reached (*t*_*max*/2_) is not easy to estimate and shows significant variability between genotypes and years. Year 2016 shows generally earlier *t*_*max*/2_ although the water stress was less severe as compared to that experienced in 2015. Consistency of the ranking between the 8 common genotypes is poor (ρ=0.33) for *t*_*max*/2_, making it a trait difficult to use for breeding. The rate of development of leaf rolling at the leaf (score) and canopy levels (CLIR) shows little differences between years, especially for CLIR that changes with similar paces for the 2 years. Although the genotypic variability is significant, the consistency of the ranking between the common genotypes for the 2 years is poor both for the rate of change of the score (ρ=0.21) and for CLIR (ρ=0.10). As a result, the magnitude of the rolling from the early morning to the maximum rolling value appears to be the main trait that was related to some genotype features. This should be further investigated on a larger scale to quantify the corresponding heritability and possible association with markers in the genome. A high-throughput phenotyping method should therefore be developed to estimate leaf rolling from canopy level measurements. The use of UAVs equipped with a multispectral camera would provide a very efficient way to cover a large experiment within a limited time period. A minimum of two flights will be necessary: one in the early morning to document the unrolled state, and one in the mid-afternoon to quantify the canopy structure when leaf rolling is at its maximum. The measurements should be completed in periods when water stress is already well expressed, and in a day with a high VPD values to maximize plant reactions. The relation between the data captured by the multispectral camera and the leaf rolling state could be achieved simply using empirical transfer functions. A representative sample of ground measurements should therefore be collected concurrently to the flights to calibrate the transfer functions. The proposed *FIPAR*_*WS*_ variable as derived from DHPs according to the methodology presented in this study would provide an efficient solution.

Although leaf rolling can be quantified both at the leaf and the canopy levels as demonstrated in this study, differences between genotypes in terms of physiological response to water stress is still a pending question. The differences between genotypes may relate to variations in soil moisture due to differences in water consumption or rooting system development and efficiency. It may also relate to the regulation of stomatal closure, as well as variation in the relation between rolling at the leaf level and leaf water potential induced by morphological leaf differences. Further, while this study shows that the leaf level rolling induces canopy level changes relatively stable across genotypes, possible residual genotypic effects may complicate the interpretation. Detailed studies are therefore required to better understand the mechanisms that sustain leaf rolling under water stress conditions.

## 5 Acknowledgement

This study was supported by “Programme d’investissement d’Avenir” PHENOME (ANR-ll-INBS-012). The experiment was mostly conducted by BIOGEMMA.

## References

Abd Allah, A.A., 2009. Genetic studies on leaf rolling and some root traits under drought conditions in rice (Oryza sativa L.). African Journal of Biotechnology 8, 6241–6248.

Adebayo, M.A., Menkir, A., 2014. Assessment of hybrids of drought tolerant maize (Zea mays L.) inbred lines for grain yield and other traits under stress managed conditions. Nigerian Journal of Genetics 28, 19–23.

Bolanos, J., Edmeades, G., 1996. The importance of the anthesis-silking interval in breeding for drought tolerance in tropical maize. Field Crops Research 48, 65–80.

Clarke, J.M., 1986. Effect of leaf rolling on leaf water loss in Trilicam spp. Canada. Journal of Plant Science 66, 885–891.

Demarez, V., Duthoit, S., Weiss, M., Baret, F., Dedieu, G., 2008. Estimation of leaf area index (LAI) of wheat, maize and sunflower crops using digital hemispherical photographs. Agricultural and Forest Meteorology 148, 644–655.

Divi, U.K., Rahman, T., Krishna, P., 2010. Brassinosteroid-mediated stress tolerance in Arabidopsis shows interactions with abscisic acid, ethylene and salicylic acid pathways. Bmc Plant Biology 10.

Driscoll, S., Prins, A., Olmos, E., Kunert, K., Foyer, C., 2006. Specification of adaxial and abaxial stomata, epidermal structure and photosynthesis to C02 enrichment in maize leaves. Journal of experimental botany 57, 381–390.

Duncan, W.G., 1971. Leaf angles, leaf area and canopy photosynthesis. Crop Science 11, 482–485.

Farhangfar, S., Bannayan, M., Khazaei, H.R., Baygi, M.M., 2015. Vulnerability assessment of wheat and maize production affected by drought and climate change. International Journal of Disaster Risk Reduction 13, 37–51.

Hay, J.O., Moulia, B., Lane, B., Freeling, M., Silk, W.K., 2000. Biomechanical analysis of the Rolled (RLD) leaf phenotype of maize. American Journal of Botany 87, 625–633.

Jonckheere, I., Fleck, S., Nackaerts, K., Muys, B., Coppin, P., Weiss, M., Baret, F., 2004. Review of methods for in situ leaf area index determination: Part I. Theories, sensors and hemispherical photography. Agricultural and Forest Meteorology 121, 19–35.

Kadioglu, A., Terzi, R., 2007. A dehydration avoidance mechanism: Leaf rolling. Botanical Review 73, 290–302.

Kadioglu, A., Terzi, R., Saruhan, N., Saglam, A., 2012. Current advances in the investigation of leaf rolling caused by biotic and abiotic stress factors. Plant Science 182, 42–48.

Krishna, P., 2003. Brassinosteroid-mediated stress responses. Journal of Plant Growth Regulation 22, 289–297.

Lopez-Lozano, R., Baret, F., Chelle, M., Rochdi, N., Espana, M., 2007. Sensitivity of gap fraction to maize architectural characteristics based on 4D model simulations. Agricultural and Forest Meteorology 143, 217-.

Monteith, J., Unsworth, M., 2007. Principles of environmental physics Third edition. Academic Press, London (UK).

Moulia, B., 1994. Biomechanics of leaf rolling. Biomimetics 2, 267–281.

Moulia, B., 2000. Leaves as shell structures: double curvature, auto-stresses, and minimal mechanical energy constraints on leaf rolling in grasses. Journal of Plant Growth Regulation 19, 19–30.

Nar, H., Saglam, A., Terzi, R., Varkonyi, Z., Kadioglu, A., 2009. Leaf rolling and photosystem II efficiency in Ctenanthe setosa exposed to drought stress. Photosynthetica 47, 429–436.

O'Toole, J.C., Cruz, R.T., Singh, T.N., 1979. Leaf rolling and transpiration. Plant Science Letters 16, 111–114.

Peleg, Z., Fahima, T., Krugman, T., Abbo, S., Yakir, D.e.a., 2009. Genomic dissection of drought resistance in durum wheat × wild emmer wheat recombinant inbreed line population. Plant Cell and Environment 32, 758–779.

Premachandra, G.S., Saneoka, H., Fujita, K., Ogata, S., 1993. Water stress and potassium fertilization in field grown maize (Zea mays L.): effects of leaf water relations and leaf rolling. J Agron Crop Sci 170, 195–201.

Price, A.H., Cairns, J.E., Horton, P., Jones, H.G., Griffiths, H., 2002. Linking drought-resistance mechanisms to drought avoidance in upland rice using a QTL approach: progress and new opportunities to integrate stomatal and mesophyll responses. Journal of experimental botany 53, 989–1004.

Saglam, A., Kadioglu, A., Demiralay, M., Terzi, R., 2014. Leaf rolling reduces photosynthetic loss in maize under severe drought. Acta Botanica Croatica 73, 315–332.

Saglam, A., Terzi, R., Nar, H., Saruhan, N., Ayaz, F.A., Kadioglu, A., 2010. Inorganic and organic solutes in apoplastic and symplastic spaces contribute to osmotic adjustment during leaf rolling in ctenanthe setosa. Acta Biologica Cracoviensia Series Botanica 52, 37–44.

Sankaran, S., Khot, L.R., Espinoza, C.Z., Jarolmasjed, S., Sathuvalli, V.R., Vandemark, G.J., Miklas, P.N., Carter, A.H., Pumphrey, M.O., Knowles, N.R., 2015. Low-altitude, high-resolution aerial imaging systems for row and field crop phenotyping: A review. European Journal of Agronomy 70, 112–123.

Sarieva, G.E., Kenzhebaeva, S.S., Lichtenthaler, H.K., 2010. Adaptation potential of photosynthesis in wheat cultivars with a capability of leaf rolling under high temperature conditions. Russian Journal of Plant Physiology 57, 28–36.

Saruhan, N., Saglam, A., Kadioglu, A., 2012. Salicylic acid pretreatment induces drought tolerance and delays leaf rolling by inducing antioxidant systems in maize genotypes. Acta Physiologiae Plantarum 34, 97–106.

Sirault, X.R.R., Condon, A.G., Wood, J.T., Farquhar, G.D., Rebetzke, G.J., 2015. “Rolled-upness“: phenotyping leaf rolling in cereals using computer vision and functional data analysis approaches. Plant Methods 11.

Smith, W.K., Vogelmann, T.C., Bell, D.T., DeLucia, E.H., Shepherd, K.A., 1997. Leaf form and photosynthesis. Bioscience 47, 785–793.

Soares, A.S., Driscoll, S.P., Olmos, E., Harbinson, J., Arrabaga, M.C., Foyer, C.H., 2008. Adaxial/abaxial specification in the regulation of photosynthesis and stomatal opening with respect to light orientation and growth with C02 enrichment in the C4 species Paspalum dilatatum. New Phytologist 177, 186–198.

Subashri, M., Robin, S., Vinod, K.K., Rajeswari, S., Mohanasundaram, K., Raveendran, T.S., 2009. Trait identification and QTL validation for reproductive stage drought resistance in rice using selective genotyping of near flowering RILs. Euphytica 166, 291–305.

Takahashi, T., Kakehi, J.-I., 2010. Polyamines: ubiquitous polycations with unique roles in growth and stress responses. Annals of Botany 105, 1–6.

Talaat, N.B., Shawky, B.T., 2012. 24-Epibrassinolide ameliorates the saline stress and improves the productivity of wheat (Triticum aestivum L.). Environmental and Experimental Botany 82, 80–88.

Tatar, O., Brueck, H., Gevrek, M.N., Asch, F., 2010. Physiological responses of two Turkish rice (Oryza sativa L.) varieties to salinity. Turkish Journal of Agriculture and Forestry 34, 451–459.

Weiss, M., Baret, F., Smith, G.J., Jonckheere, I., Coppin, P., 2004a. Review of methods for in situ leaf area index (LAI) determination: Part II. Estimation of LAI, errors and sampling. Agricultural and Forest Meteorology 121, 37–53.

Weiss, M., Baret, F., Smith, G.J., Jonckheered, I., Coppin, P., 2004b. Review of methods for in situ leaf area index determination, part II: Estimation of LAI, errors and sampling. Agricultural and Forest Meteorology 121, 37–53.

